# Visual delay affects force scaling and weight perception when lifting objects in virtual reality

**DOI:** 10.1101/504563

**Authors:** Vonne van Polanen, Robert Tibold, Atsuo Nuruki, Marco Davare

**Affiliations:** Movement Control and Neuroplasticity Research Group, Biomedical Sciences, KU Leuven, Leuven 3001, Belgium; Sobell Department of Motor Neuroscience and Movement Disorders, Institute of Neurology, University College London, London WC1N3BG, UK; Central for General Education, Kagoshima University, 1-21-30, Korimoto, Kagoshima, 890-0065, Japan

**Keywords:** Grasping, multisensory, virtual reality, weight perception, force

## Abstract

Lifting an object requires precise scaling of fingertip forces based on a prediction of object weight. At object contact, a series of tactile and visual events arise that need to be rapidly processed online to fine-tune the planned motor commands for lifting the object. The brain mechanisms underlying multisensory integration serially at transient sensorimotor events, a general feature of actions requiring hand-object interactions, are not yet understood. Here we tested the relative weighting between haptic and visual signals when they are integrated online into the motor command. We used a new virtual reality setup to desynchronize visual feedback from haptics, which allowed us to probe the relative contribution of haptics and vision in driving participants’ movements when they grasped virtual objects simulated by two force-feedback robots. We found that visual delay changed the profile of fingertip force generation and led participants to perceive objects as heavier than when lifts were performed without visual delay. We further modeled the effect of vision on motor output by manipulating the extent to which delayed visual events could bias the force profile, which allowed us to determine the specific weighting the brain assigns to haptics and vision. Our results show for the first time how visuo-haptic integration is processed at discrete sensorimotor events for controlling object lifting dynamics and further highlight the organization of multisensory signals online for controlling action and perception.

**New & Noteworthy:** Dexterous hand movements require rapid integration of information from different senses, in particular touch and vision, at different key time points as movement unfolds. The relative weighting of vision and haptics for object manipulation is unknown. We used object lifting in virtual reality to desynchronize visual and haptic feedback and find out their relative weightings. Our findings shed light on how rapid multisensory integration is processed over a series of discrete sensorimotor control points.

## Introduction

Skilled object lifting involves the planning of a motor command specifying fingertip forces based on a prediction of the object weight. In case of a mismatch between the predicted and actual sensory feedback, forces are rapidly adjusted to control the action (Johansson and Westling 1988). When lifting an object, multiple sources of sensory inputs are available to adjust fingertip forces to the specific object properties. For instance, haptic information about object weight and material is retrieved from both proprioceptive sensors as well as mechanoreceptors in the fingertips, just after object contact. Decreased grip force control is seen in the absence of cutaneous information (e.g. by anaesthesia, Johansson and Westling 1984; Monzée et al. 2003), or force feedback (Gibo et al. 2014). In addition, visual events are also available in terms of object movement during lifting and visible contact of the fingertips with the object that indicate a stable grasp. Absence of visual information leads to impaired adaptation of force scaling in repeated object lifting (Buckingham and Goodale 2010). Furthermore, observing others handling objects can influence subsequent force scaling as well (Buckingham et al. 2014; Uçar and Wenderoth 2012).

Multisensory integration has been widely studied in the context of perception. That is, how the brain generates a perceptual estimate of a given object property depending on available sources of sensory information. This definition distinguishes between actual sensory signals, which is the input processed in the brain, and perception, which is the final estimation of an object property that can be reported by a subject. In general, combined sensory input provides a better percept than when only one sensory modality is available by optimal integration of the unimodal information sources (Ernst and Banks 2002; Helbig and Ernst 2007). The optimal integration is also evident in findings that the weighting assigned to modalities depends on the accuracy of the information carried by a given sensory channel (Ernst and Banks 2002; Helbig and Ernst 2007; Knill and Saunders 2003).

Neural mechanisms underlying multisensory integration for controlling actions might be different than those underlying perception. On the one hand, when generating a perceptual estimate of a given object property, the brain usually receives constant sensory inputs in parallel from different sources and for a substantial amount of time. On the other hand, for action control, sensory information is rather processed serially at discrete sensorimotor control points even though multisensory signals are available continuously (Johansson and Flanagan 2009). For example, during object lifting, key visual time points are initial contact of the fingers with the object and object movement (lift-off). In addition, these events are sensed in parallel by fast adapting tactile mechanoreceptors in the skin (Johansson and Flanagan 2009). These sensorimotor control points are critical because the brain must quickly adjust the ongoing motor command if the actual sensory feedback from either tactile or visual signals at these time points deviates from expected values. Therefore, the rapid succession of these sensorimotor events and the rapid motor response they require make it very likely that sensory feedback from vision and touch is processed differently for action control than for perception. For instance, the relative weighting of visual and haptic inputs might be different for controlling actions than for generating perceptual estimates.

In this study, we wanted to investigate the relative contribution of visual and haptic information to controlling object lifting. Since visual and haptic signals are intrinsically not dissociable in real life, i.e. they provide the same information (contact, lift-off) at these different control points, such a question is merely impossible to investigate with real objects. Therefore, we used a virtual reality environment to introduce a visual delay with respect to haptic signals while subjects lifted virtual objects. By desynchronizing visual from haptic signals, our goal was to estimate the relative weighting of vision and haptics by quantifying how much visual delay could affect force generation. That is, the larger the effect of visual delay, the larger the gain of vision relative to haptics.

Moreover, we recently showed that the lifting phase is important for mediating weight perception (van Polanen and Davare 2015). Naturally, lifting an object provides information about its weight. Therefore, a second aim of this study was to determine whether visual delay would also affect perceptual weight judgments of objects that are actively lifted. It is worth mentioning that no study investigated the effect of visuo-haptic asynchronies on both force control and perception of weight. Hence, it is undetermined whether the contribution of vision to these processes is related or not, which would provide valuable information about how perception and action processes interact.

It is noteworthy that previous studies investigating visual delay effects on object manipulation only considered the object holding phase. For instance, when holding or transporting objects, an altered grip and load force coupling (Sarlegna et al. 2010) and increased weight perception (Honda et al. 2013; Kambara et al. 2013) were found, suggesting a role of vision in force control and weight perception. Critically, these studies did not examine object lifting and therefore do not address the mechanisms underlying multisensory integration for implementing rapid sensorimotor corrections based on a mismatch between expected and actual sensory feedback. This mismatch can be detected at each sensorimotor control time point via visual and haptic feedback, and between object contact and lift-off. Therefore, current knowledge about the relative contribution of visual and tactile events for controlling force generation via rapid feedback loops is still lacking.

We addressed these issues in two experiments, in which objects were lifted with and without visual delays and participants judged the heaviness of the lifted objects. We found that a visual delay alters force scaling and also increases perceived object weight. These two effects were not strongly related. To determine the relative weighting of visual and haptic information, we compared different models each explaining different mechanisms combining multimodal inputs for motor control. We found that visual and haptic information contribute rather independently to force generation.

## Methods

### Participants

Thirty participants (15 female, two left-handed) took part in the study, of which 14 participated in Experiment 1 (30.14±4.05 years) and sixteen in Experiment 2 (24±4.5 years). Participants provided informed consent before the experiment and had no known visual deficits (including lack of stereovision). After debriefing, only one subject (Experiment 1) reported she could notice the delay without the ability to quantify its duration. The experiment was approved by the local ethical committee of UCL (Experiment 1) and KU Leuven (Experiment 2).

### Apparatus and task

We used a virtual reality setup with two phantom haptic devices (Sensable), as illustrated in Figure 1. A projected visual 3D scene in combination with the haptic devices provided a realistic grasping and lifting experience. Participants were seated in front of a table, with their thumb and index fingertips inserted into the thimbles of each phantom. These devices were placed underneath a mirror, thus the participant’s hand and haptic devices were not visible. The mirror projected a 3D-screen (Experiment 1: LG, Experiment 2: Zalman) to provide a visual virtual environment that was aligned with the actual finger positions of the participant. The virtual environment consisted of a background with black and white squares to provide perspective cues. In addition, the fingertip positions (red spheres), the lifted object (blue cube), two start positions (red-green poles, one for each finger, not shown in Figure 1) and a target location (yellow mark) were present. The object and fingertip positions were continuously visually displayed during a trial. In response to object contact and squeeze, the red spheres (fingertips) could slightly penetrate the object surface, such that only part of the sphere became visible. This mimics fingertip deformation when squeezing an object in real-life conditions and enhanced the impression of squeezing a veridical object in our virtual reality setup.

**Figure 1.**
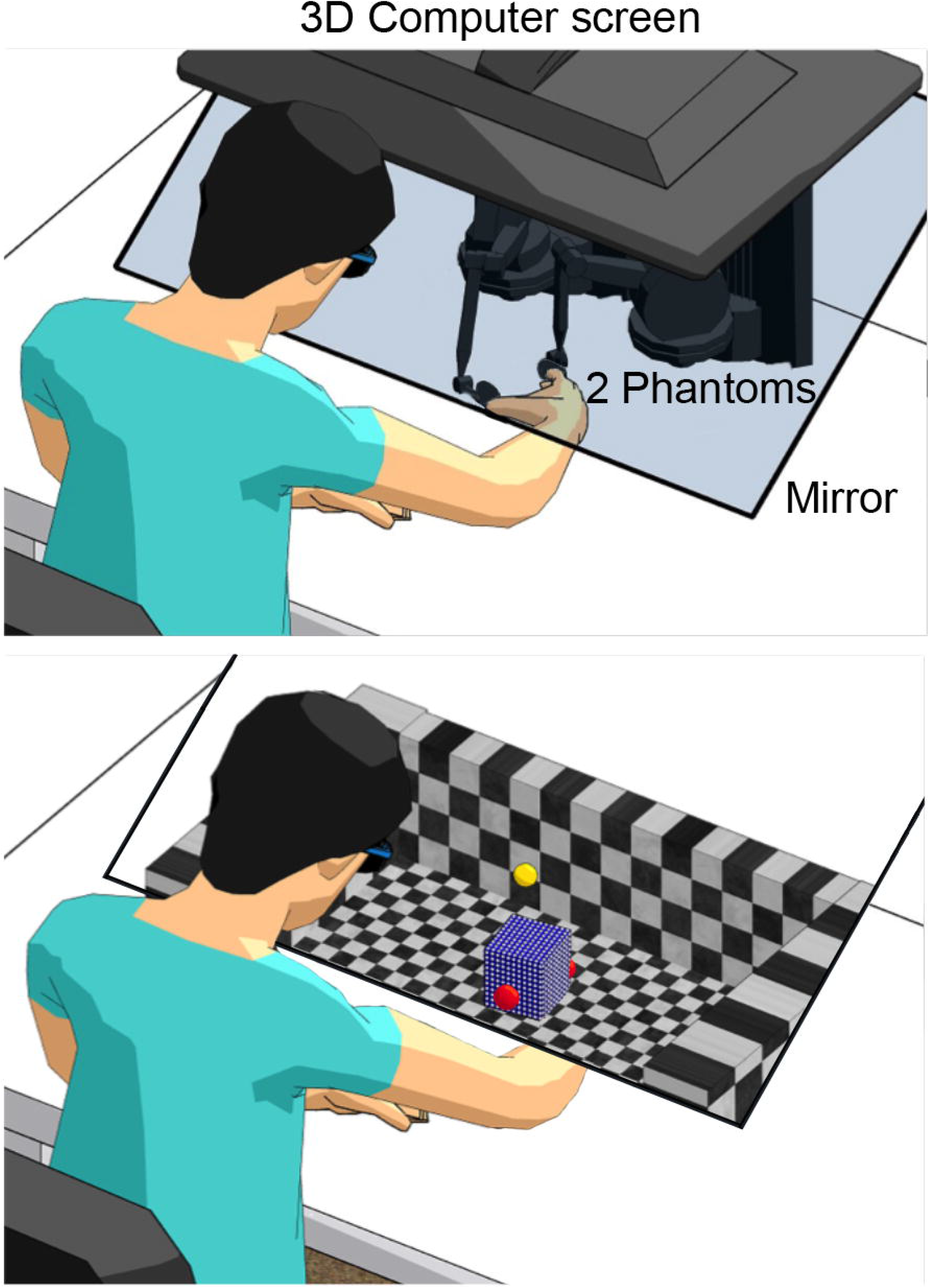
Experimental set-up (top) and virtual reality environment (bottom). The red spheres represent the participant’s fingertips. The blue cube had to be lifted up to a height indicated by the yellow mark.

The haptic devices provided force feedback to the participant and measured position and force information in three directions, sampled at 500 Hz. In response to position and velocity changes of the haptic devices, the position and acceleration of the object was calculated and the resulting forces that were applied to the participant’s fingers were determined. Therefore, generated grip and load forces automatically followed the movement of the participant. Normal forces applied to the cube (grip forces) were modeled as a spring, where the stiffness of the box was set at 0.4 N/mm. This low stiffness value was chosen to avoid overload of the haptic device when high opposing forces need to be generated. Vertical (load) forces were the summation of the gravitational (g=9.81 m/s^2^), angular momentum and damping forces (damping constant is 2 kg/s). For force calculations, the openHaptics toolkit was used, which sends commands and receives information from the devices at a rate of 1kHz. Force commands were sent to the devices within a single sample to synchronize the devices with minimal delay (i.e. maximum 1 ms). The visual delay due to the screen refresh rate (60Hz) was 17 ms. These intrinsic delays, due to the virtual reality setup, were present in all conditions and treated as a baseline value (i.e. 0 ms experimentally induced delay).

Participants were positioned in front of the set-up and familiarized with the procedure. They performed practice trials to get used to the virtual reality environment. In the experimental trials, a virtual cube was positioned in the center of the environment and participants were instructed to hold their fingertips at the virtual start positions (i.e. one fingertip at each position) that were shown in front of the participant, between the participant and the cube. After a beep (1.5 s after the cube appeared), they could reach for the cube, grasp it and lift it up to the target level (Figure 1). Participants were instructed to hold the cube up to this level until a second beep (3.0 s after the first beep). Afterwards they had to replace the cube back on the virtual table. The cube disappeared and participants waited until the experimenter initiated the next trial. The visual appearance and size of the cube was always the same such as not to give any cues about its mass.

### Experiment 1: Lifting objects of different masses with or without delay

In Experiment 1, participants lifted cubes of different masses that could be presented with or without a delay. In the case of a delay, the delay was present during the whole time the cube was visible, i.e. during the complete trial. Participants estimated the heaviness of the cube after each lift by providing a number best representing the perceived heaviness on a self-chosen scale (magnitude estimation, Zwislocki and Goodman 1980). Four cube masses were used (100, 200, 300 and 400 g) and two delays (100 and 200 ms). The order of trials was randomized, with 10 trials for each combination of cube mass and delay (total of 80). In addition, all masses were also presented without a visual delay. To make the delay less noticeable, two times as many trials without delay (40 for each mass, total 160) than with delay were presented. A total of 240 trials were performed.

### Experiment 2: Comparing two cubes lifted with and without delay

In order to better quantify the effect of visual delay on weight perception, we performed a staircase procedure to determine the weight perception bias caused by a visual delay. In this experiment, participants lifted two visually identical cubes and indicated which of the two cubes felt heavier. The cubes were presented sequentially, similar to two consecutive trials in Experiment 1, and each cube was only lifted once. When lifting a cube with a visual delay, the delay was present the entire time the cube was visible. After lifting the second cube, participants reported whether the first or second cube felt heavier. Since there was limited time between the presentations of the two cubes, so that first object did not need to be kept in memory for a long time, participants were not instructed to return to the start positions, but could choose their own preferred starting position in this experiment.

Critically, in one of the two compared lifts, visual information was delayed with respect to haptics by either 100 or 200 ms, whereas in the other lift no delay was present. In this way, lifts with and without delay were directly compared. An adaptive staircase procedure (one up one down) was followed where a standard mass of 200 g was compared with a variable test mass (ranging between 110-290 g). Two staircases were interleaved with test masses starting at 110 or 290 g. After each comparison, the next test cube was increased with 15 g when it was perceived as lighter, or decreased if it was perceived as being heavier. Staircases were terminated after 15 comparisons (30 lifts in each session), which was enough to reach stable performance (except for one participant in one session, see below).

Eight participants took part in the 100 ms delay condition and the other eight performed the 200 ms delay condition. In each delay group, two sessions were performed. In one session the standard mass was delayed and in the other session the test mass was delayed. This allowed us to determine which non-delayed test mass was perceived to be equally heavy as the delayed standard mass or, in the other session, which delayed test mass felt the same as the non-delayed standard mass. Session order was counterbalanced among participants and the order of standard and test mass presentation (and delayed or non-delayed cubes) was randomized.

### Analysis of perceptual estimates and biases

In Experiment 1, participants’ answers were converted to z-scores and averaged over cube mass and visual delay. In Experiment 2, we calculated the perceptual bias, to determine whether a cube lifted with a visual delay was perceived differently compared to a non-delayed cube. The percentage of answers when the test mass was reported ‘heavier’ were calculated for each presented comparison, for the two sessions separately. The percentages were plotted and a psychometrical curve was fitted to the points:

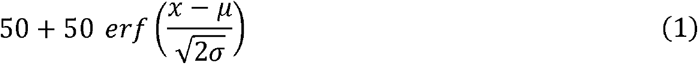

where *μ* and *σ* are the fitted parameters representing the mean and standard deviation of the curve, respectively. Because some test masses were presented more often than others, a weighted fit was used. The value of *μ* represents the perceptual bias for a specific session. The total bias was calculated by subtracting the bias for the test-delay session from that of the standard-delay session, dividing this by 2 and presented as a percentage difference compared to the standard weight of 200 g. This percentage indicates how much heavier (positive value) or lighter (negative value) a cube lifted with a delay is perceived compared to a cube lifted without a delay.

For two participants in the 200 ms delay experiment, in one of the sessions the bias was bigger than the difference between the standard and the maximum test mass. In those cases, the bias was set to the maximum possible value in our set (i.e. 45%). In one participant in one session, the staircases did not converge completely before the 15 comparisons were reached. This participant was very variable in reporting which cube was heavier, possibly due to low weight sensitivity or strategy differences for heaviness determination. However, since only one test mass could not be compared to the standard for this participant we decided to still include data from this participant in the analysis and just exclude this missing test value from the fit.

### Analysis of force scaling parameters

To investigate the effect of visual delay on force scaling, several force parameters were calculated. Forces were filtered with a 2^nd^-order bidirectional low-pass Butterworth filter, with a cut-off frequency of 15 Hz. Missing samples (<0.1%) were linearly interpolated. Trials with multiple lifts, dropped cubes during lifting or technical errors were excluded from analysis (3%). Parameters were defined for each cube mass (100-400 g) in Experiment 1. To exclude effects of cube mass on force scaling in Experiment 2, only lifts for the 200 g cubes (standard mass) were analyzed. The differences between forces in delay and no-delay conditions were further quantified by calculating parameters indicative of force scaling. The four parameters of interest were the maximum load force rate (LFRmax), maximum grip force rate (GFRmax), loading phase duration (LPD) and the grip force at the time of lift-off (GFatLO). Load force (LF) was defined as the sum of the two vertical components (tangential to the object surface) of the forces applied by the thumb and index fingertips. Grip force (GF) was the mean of the horizontal (perpendicular to the object surface) force components. LF onset and GF onset were set at the time point forces reached a threshold of 0.1 N. Lift-off was determined as the time point when LF became equal to the weight of the lifted cube. Load force rates (LFR) and grip force rates (GFR) were the differentiated LF and GF, respectively. Maximum LFR (LFRmax) and GFR (GFRmax) were calculated as the maximum force rate between 50 ms before onset and 50 ms after lift-off. A slightly larger time range was used for two reasons: 1) since onset was determined based on forces, not force rates, peaks could already appear early (before onset, <0.1% trials) and 2) especially if lift-off occurs early than expected, force peaks can also occur just after lift-off (1.9% trials). The load phase duration (LPD) was the time between LF onset and lift-off.

Since the force parameters only give information about a specific time point during the lifting movement, we also performed an exploratory analysis on the force profiles. To evaluate the difference in the force profiles in delay and no-delay lifts, we calculated the average force and force rate curves for each participant and condition. Force and force rate curves were aligned at GF onset and analyzed starting from this time point onwards for a period of 1s, by bins of 100 ms (10 bins). The area under the curve was calculated for each cube mass, delay and bin.

We explored the relation between force parameters and perceptual biases by calculating correlation coefficients across participants. In this way, we could compare the results of processing of visual and haptic signals for weight perception and force control. Relative differences between delay and no-delay conditions were determined for 100 and 200 ms delay separately. These relative changes in force parameters were correlated with the relative z-score changes (Experiment 1) or perceptual biases (Experiment 2). Correlations were performed for each delay, cube mass and force parameter, giving 32 and 8 correlations in Experiment 1 and 2, respectively. Positive correlations indicate that force parameter values as well as heaviness perception increase in response to a visual delay.

Kinematic measures were analyzed to determine whether a visual delay affected the reaching phase before contact. This analysis was only performed in Experiment 1, because participants did not always start from the fingertip start positions in Experiment 2, almost completely removing the reaching phase. Trials in which participants were not at the start position at the beginning of the trial were removed for this analysis (2%). Kinematic parameters were determined from the start of the movement (defined as the moment one of the fingers reached a velocity of 10 mm/s in the direction towards the cube) until object contact, as indicated by GF onset. The investigated parameters were peak velocity, travelled path, path curvature (maximum deviation from a straight line) and were averaged over the two fingers and collapsed over cube mass. In addition, the position at contact in the reach direction was determined for the fingers separately. The path curvature and position at contact were analyzed to further investigate the differences we found in the travelled path (see Results).

### Statistics

Statistical analyses were performed with SPSS version 24 (IBM). In Experiment 1, variables were analyzed with a 3 (delay) × 4 (mass) repeated measures Analysis of Variance (ANOVA). Variables of interest were the z-scored weight percept values, LFRmax, GFRmax, LPD and GFatLO. Since kinematic variables (peak velocity, travelled path, path curvature and finger position) were collapsed over cube mass, they were analyzed with an ANOVA with a single factor (delay, 3 levels). Post-hoc tests were performed with paired samples *t*-tests with a Bonferroni correction. If the sphericity assumption was violated (Mauchly’s test), a Greenhouse-Geisser correction was applied.

For the analysis of the force profiles, we were interested in how the delay conditions differed at different time points. Since effects of mass and bin would be trivial, we performed planned comparisons on the effects of delay. We compared the three conditions of delay: no-delay, 100 ms and 200 ms delay. Conditions were compared (3 comparisons for each bin) using paired *t*-tests with a Bonferroni correction to account for the 10 bin comparisons. Comparisons were made for each cube mass separately.

In Experiment 2, a 2 × 2 ANOVA was used with the within factor delay (delay or no-delay) and the between factor group (100 ms, 200 ms). The analyzed variables were LFRmax, GFRmax, LPD and GFatLO. One-sample *t*-tests were used to determine whether the total biases were significantly different from zero (*p* < 0.05).

### Modeling of load force curves

To quantify the relative weighting of visual and haptic information on force scaling, we modeled the force planning in response to a visual delay. Modeling complex corrective feedback mechanisms during lifting was beyond the scope of this paper. Instead, we used simple models that incorporate haptic and visual feedback about object contact and tested how these affected the force generation profile. We considered three different models in which visual delay could affect the force profile (Figure 2A): (1) the force generation profile was kept unchanged but its onset was shifted by visual delay (shift model), (2) visual delay slowed down the force generation profile, but its onset was unaltered (stretch model) and (3) a combination of a ‘haptic’ (i.e. non-shifted) and ‘visual’ (i.e. shifted) force profile (sum model). In all models, the relative weight given to visual information determined how much the original force curve without delay was altered by visual delay. If one completely relies on vision (weight=1), the curve is maximally altered. On the other hand, if one completely relies on haptic information (weight=0), no change in force scaling is expected with visual delay. The models were calculated using Matlab 2017 (MathWorks). The program code can be found in the Supplemental Material.

**Figure 2.**
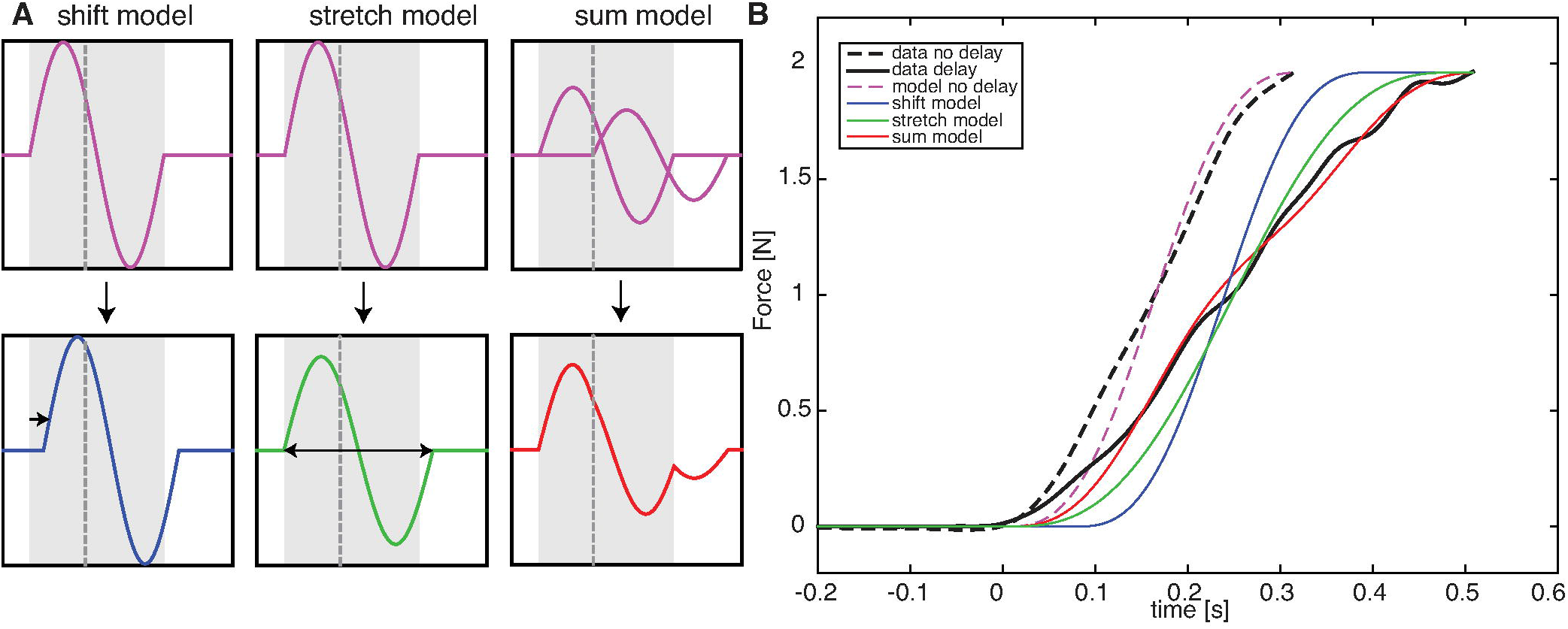
A. Illustration of the three models (shift, stretch and sum). Shaded areas are drawn between GF onset and original lift-off, dashed lines indicate the visual delay (e.g. 100 or 200 ms). For each model in this example, w_h_=0.75 and w_v_=0.25. The second derivative of the load force is modeled by a basic sine function (purple). In the shift model (blue), the basic sine is shifted between haptic contact and visual contact. In this case, the shift is 25% of the visual delay. In the stretch model (green), the duration of the sine is increased with, in this case, 25% of the visual delay duration. The sum model (red) is a combination of a sine at haptic contact and at visual contact. The amplitude of both sines is determined by the weighting factor and is, in this case, 75% of the basic sine for the haptic curve and 25% for the visual curve. B. Example of three fitted models for a single participant lifting a 200 g cube. Black traces represent averaged actual data plotted until lift-off. Dashed lines are no-delay trials, solid lines are 200 ms delay trials. For this example, w_h_ is 0.62, 0.16 and 0.59 for the shift, stretch and sum models, respectively.

The force curves were modeled as a sigmoid curve starting to rise at GF onset, corresponding to object contact, and saturating at lift-off at a value equal to the cube’s weight. It is appropriate to apply sigmoid fitting methods to fingertip force output during object lifting. When correctly scaled to the object weight, fingertip force rates follow a bell-shaped curve and are symmetrical. Since we were interested in modeling the loading phase, the force curve model was calculated until lift-off. Because at lift-off the load forces have to be equal to cube weight, it was valid to assume that the model’s final force value would be the same as the cube’s weight. A sigmoid curve is the double integration of a sine with a specific frequency and amplitude of a single period. By altering the frequency, onset and amplitude of this force acceleration, we could define the final force curve in different ways: the onset defines at what time point the force starts to increase (e.g. at object contact), the amplitude and frequency determine at what value the sigmoid ends (e.g. cube weight) and how long it takes to reach this level (e.g. loading phase duration). The starting point was chosen at GF onset, which represents object contact and is a good approximation of the expected LF acceleration onset. Due to noise in the second derivative of LF signals, GF onset was more reliably determined than LF acceleration. The resulting sigmoid function was fitted to the average LF curves from the data for each participant and each condition.

#### Basic model: sine wave

The basic force acceleration was defined as a sine wave of the form *G* sin (*ft*), where *G* is the amplitude of the sine, *f* the frequency and *t* the independent variable time. The amplitude determines the final force value that is reached and the time to reach this force is specified by the frequency. Because the final force value should be equal to the cube weight (*m*), only the time to reach this value (here called duration, *d*) is a free parameter. Therefore:

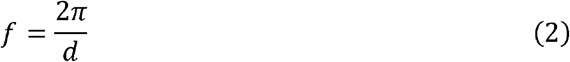

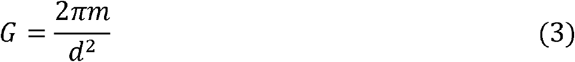

We defined three models that could be described as a shifted sine (shift model), a stretched sine (stretch model) or a weighted combination of two sines (sum model). In each model, only *d* and the relative weight of vision (*w_v_*) and haptics (*w_h_*) were the two free parameters. Because *w_v_* and *w_h_* sum up to 1, these weights were considered as a single free parameter. The models are visualized in Figure 2.

#### Model fitting of no delay trials (duration d) and delay trials (weights w_v_ and w_h_)

In all models, it was assumed that *d* was independent of the delay, but specific for each participant and each cube mass. Therefore, it was assumed that participants planned the same lift duration irrespective of a delay. The no-delay trials were used to obtain a measure of *d* for each participant and cube mass. Since in no-delay trials visual and haptic information are synchronous, they provide the same information and their weighting is redundant. The only remaining unknown parameter *d* is found by fitting one of the models to the averaged no-delay LF curve (note that all models give the same result without a delay, see black lines in Figure 2B and grey triangles in Figure 6). We fitted to average curves to make sure the force curves of the data were accurately scaled to the cube’s weight and not influenced by sensorimotor memory of the previous lift (Johansson and Westling 1988). The time point of lift-off was defined for the averaged LF curve and the modeled curve was compared with the measured curve until this time point. The fitting procedure minimized the root mean square error (RMSE):

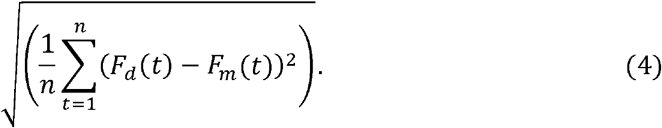

Here, *F_d_* is the force curve from the data, *F_m_* is the modeled curve and *n* is the number of data points until lift-off. In the delay conditions, the *d* from the no-delay condition was taken and the same fitting procedure was followed to define the weighting factor *w_v_* for each model by minimizing the RMSE. This value also gives a measure of goodness of fit of the model, with lower values indicating that the model fits the data better. To decide which of the three models fitted the data best, the RMSE values were compared. For Experiment 1, a repeated measures 3 (model) × 2 (delay) × 4 (mass) ANOVA was used and for Experiment 2, a repeated measures ANOVAs with a within factor (model, three levels) and a between factor delay (100 ms, 200 ms) was performed.

#### Shift model

The shift model only shifts the onset of the acceleration sine wave with respect to GF onset (i.e. object contact). This model assumes that delayed visual information will also delay the onset of force scaling. How much the curve is shifted depends on the weight given to vision (*w_v_*) and visual delay. When *w_v_*=0, the curve is not shifted and the onset remains at GF onset. When *w_v_*=1, the curve is maximally shifted with the delay. For 0<*w_v_*<1, the shift is between 0 and the visual delay value (100 or 200ms).

#### Stretch model

The stretch model only changes the duration (*d*). This means that in this model, the lift-off time point is delayed based on the visual delay. With a maximal value of *w_v_* =1, *d* increases by an amount equal to the visual delay value. For 0<*w_v_*<1, *d* is increased with a value between 0 and the delay. According to Eq. 3, the amplitude *G* depends on *d*, indicating that the amplitude will decrease with a larger *d*.

#### Sum model

In the sum model, the acceleration is calculated as a combination of two sine waves, one for each modality (Figure 2A, top right). The ‘haptic’ wave is a sine with an onset at GF onset. The ‘visual’ wave is the same sine wave, but the onset is shifted by the visual delay value. The final force acceleration is a weighted sum of both sines (Figure 2A, bottom right): the amplitude of each sine is multiplied by the weighting for the specific modality. Thus, for the haptic sine wave, the amplitude is *w_h_G* and for the visual sine wave, the amplitude is *w_v_G*. This model assumes a rather independent contribution of haptic and visual information to the force scaling. The weight of each modality determines its contribution. Note that if *w_v_* =1, only the ‘visual’ sine is used and therefore is equal to a completely shifted curve (i.e. similar to shift model with *w_v_* =1).

## Results

### Experiment 1: Visual delay alters force scaling and weight perception

We used a virtual reality environment where subjects had to lift virtual cubes and estimate their heaviness. Crucially, in some trials vision was delayed with respect to haptics by 100 or 200 ms. We found that visual delay altered the force scaling of fingertip forces applied to the cube as seen in the average force traces in Figure 3. The top row shows grip (red shades) and load forces (blue shades). The force profiles were compared within 10 bins of 100 ms. Significant differences can be seen between force profiles in delay vs. no-delay lifts and are indicated with colored bars. Specifically, load forces were lower during the loading phase, whereas grip forces were higher after liftoff. Moreover, it appears that force profiles are somewhat shifted with delay. When analyzing differences in force rates for different time bins (bottom row of Figure 3), it was found that, for the first bins, force rates were lower in the delay conditions, but for later bins, force rates were higher in the delay conditions.

**Figure 3.**
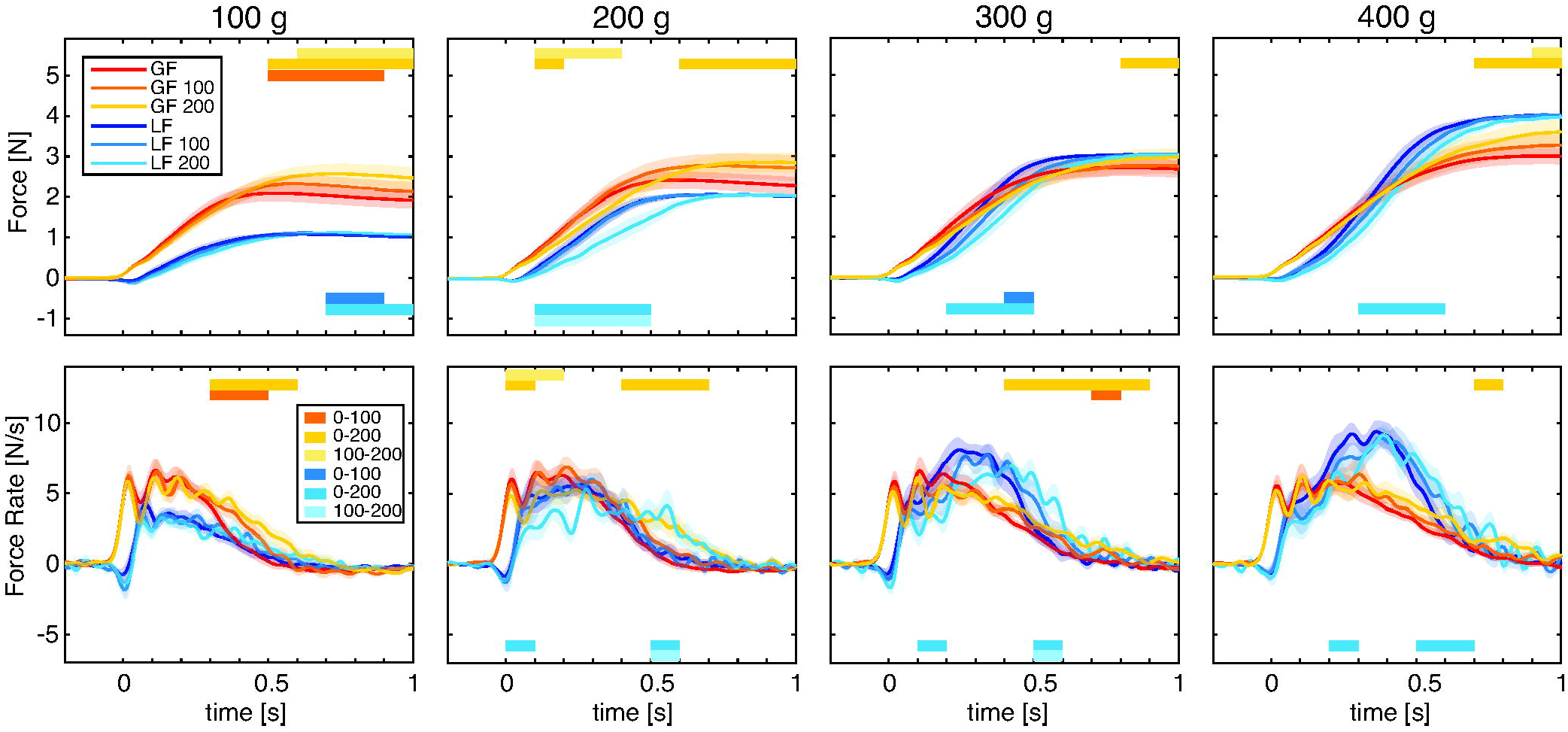
Averaged force curves (top) and force rate curves (bottom) for each cube mass and delay from Experiment 1. Traces are aligned at grip force onset (t=0). Lighter colors represent larger delays with red shades for grip forces and blue shades for load forces. Horizontal bars indicate time bins for which significant differences between delay conditions were found (paired samples t-tests between delay conditions with Bonferroni correction for the 10 bin comparisons). Transparent shades represent standard errors. Note negative load forces just after contact are due to subjects slightly pressing the virtual object on the virtual table before initiating the lift, as in real-life conditions (e.g. see van Polanen and Davare, 2015).

Overall, this suggests that visual delay led to a slower force generation and increased grip forces. We investigated force parameters indicative of force scaling shown in Figure 4. It seems that most force parameters, except GFRmax, increased with cube mass. In addition, pronounced effects of a visual delay were seen as well. For LPD, a 3 (mass) × 4 (delay) repeated measures ANOVA revealed main effects of mass (*F*_(1.6,20.5)_=148.8, *p*<0.001, *η_p_*^2^=0.92) and delay (*F*_(1.3,17.4)_=13.1, *p*=0.001, *η_p_*^2^=17.4). LPD increased with cube mass and all comparisons were significantly different. In addition, the LPD increased with delay, where all conditions differed significantly.

**Figure 4.**
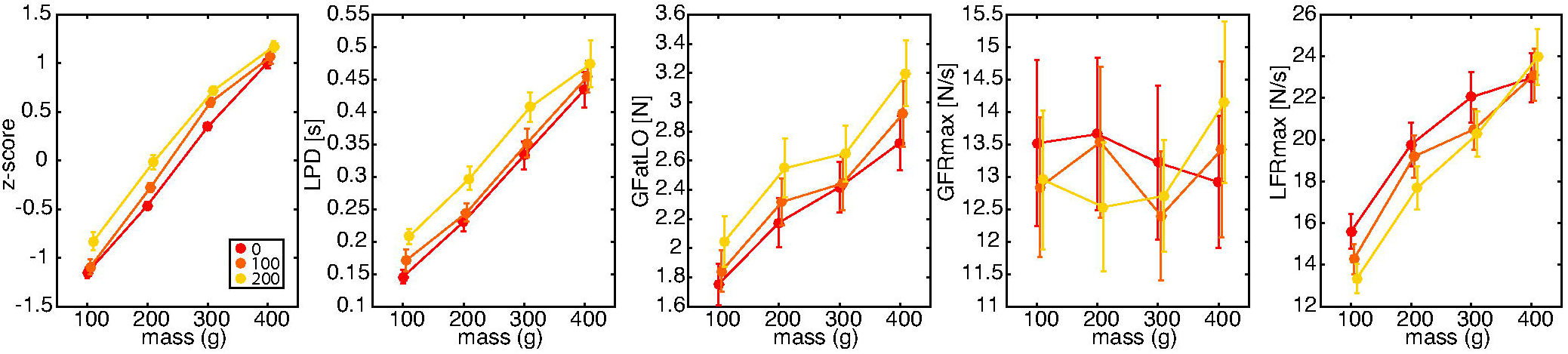
Results for Experiment 1. From left to right: magnitude estimations (z-scores), load phase duration (LPD), grip force at lift-off (GFatLO), maximum grip force rate (GFRmax) and load force rate (LFRmax). Lighter colors indicate larger delays. Error bars represent standard errors. Note that all parameters but GFRmax were significantly affected by the delay (main effect of delay in repeated measures ANOVA).

Similarly, for GFatLO effects of mass (*F*_(1.2,16.1)_=66.0, *p*<0.001, *η_p_*^2^=0.84) and delay (*F*_(2,26)_=23.1, *p*<0.001, *η_p_*^2^=0.64) were found. All mass comparisons were different, where GFatLO increased with cube mass. Furthermore, GFatLO was larger for a delay of 200 ms compared to both no-delay and 100 ms delay conditions. There was no difference between the 100 ms delay and no-delay conditions.

Although the force profiles suggest some decreases in grip force rate with a visual delay, no significant effects for mass or delay were found for GFRmax. For LFRmax, an effect of mass (*F*_(3,39)_=89.8, *p*<0.001, *η_p_*^2^=0.87), an effect of delay (*F*_(1.3,17.0)_=4.7, *p*=0.037, *η_p_*^2^=0.27) and an interaction between mass × delay (*F*_(6,78)_=5.1, *p*<0.001, *η_p_*^2^=0.28) were found. Post-hoc tests indicated that LFRmax increased significantly with cube mass, but there was no difference between the 300 and 400 g cube. Neither were the 200 and 300 g cubes different in the 100 ms and 200 ms delay conditions. The effect of delay was only significant for the 100 g cube indicating a smaller LFRmax in the 200 ms condition compared to the no delay condition.

As seen in Figure 2, a visual delay also affected magnitude estimations for lifted cubes. Higher z-scores were seen for heavier cubes, but also for lifts with larger visual delays. The ANOVA on the z-scores of the magnitude estimations revealed effects of mass (*F*_(1.5,19.0)_=265.5, *p*<0.001, *η_p_*^2^=0.95), delay (*F*_(2,26)_=36.2, *p*<0.001, *η_p_*^2^=0.74) and an interaction (*F*_(6,78)_=4.0, *p*=0.001, *η_p_*^2^=0.24). As expected, magnitude estimations increased significantly with cube mass. However, cubes were also perceived as heavier with more visual delay. For all cube masses, the 200 ms delay made cubes being perceived as heavier than cubes lifted without a delay. In addition, there was a difference between the 100 ms and 200 ms delay conditions for the 100 g cube and a difference between no and 100 ms delay conditions for the 300 g cube.

In summary, visual delay makes cubes feel heavier while it also increases grip forces and load phase durations, and decreases load force rates. To determine whether perceptual differences and effects on force scaling were related, the relative difference between delay and no-delay conditions of these parameters were correlated. Overall, from the 32 performed correlations, few correlations, and in different directions with respect to the main results, were found. We found one significant correlation of magnitude estimations with GFRmax (100 ms, 200 g cube, *R*=0.74, *p*=0.002), one with LPD (100 ms, 400 g, *R*=−0.67, *p*=0.009) and two with GFatLO (100 ms, 400 g, *R*=0.61, *p*=0.020 and 200 ms, 300 g, *R*=0.55, *p*=0.041). The first two correlations indicate that an increase in GFRmax or a decrease in LPD is associated with an increase in weight rating. These relations are surprising, since there was no significant increase in GFRmax and a significant increase in LPD was actually found, thus opposite to these correlations. The correlations with GFatLO revealed that a larger increase in grip force at lift-off was associated with a larger increase in weight perception.

Kinematic analyses indicated significant effects of visual delay on the reaching phase. Repeated measures ANOVAs with a single factor delay indicated that the fingers moved slower (lower peak velocity, *F*_(2,26)_=45.4, *p*<0.001, *η_p_*^2^=0.78) and travelled a longer path (*F*_(2,26)_=13.8, *p*<0.001, *η_p_*^2^=0.52). The longer path seems to result from a larger curvature (larger maximum deviation from straight line, *F*_(2,26)_=12.1, *p*<0.001, *η_p_*^2^=0.48) and a small overshoot (further position at contact for index finger, *F*_(1.1,14.2)_=14.6, *p*=0.002, *η_p_*^2^=0.53, and thumb, *F*_(1.2,15.9)_=15.0, *p*=0.001, *η_p_*^2^=0.54). Despite these kinematic effects on the reaching phase, subjects did not notice the visual delay, nor compensated for it later in the trial.

### Experiment 2: Quantifying the effect of visual delay on perceptual weight biases

The results of Experiment 1 showed that visual delay altered force scaling and weight perception. To further quantify the perceptual difference due to a delay, we performed a second experiment where we let participants directly compare cubes with and without a visual delay. With the use of an adaptive staircase, the perceptual bias induced by a visual delay of 100 or 200 ms was determined.

Biases are shown in Figure 5. With a delay of 100 ms, the biases were small and scattered around zero. The mean bias of 4.0±3.8% was not significantly different from zero (*t*_(7)_=1.05, *p*=0.327). On the other hand, when vision was delayed by 200 ms, most biases were positive, indicating that delayed cubes feel heavier. On average, a significant bias of 17.8±6.5% was found (*t*_(7)_=2.74, *p*=0.029). This implies that cubes of 200 g feel almost 36 g heavier when lifted with a visual delay of 200 ms.

**Figure 5.**
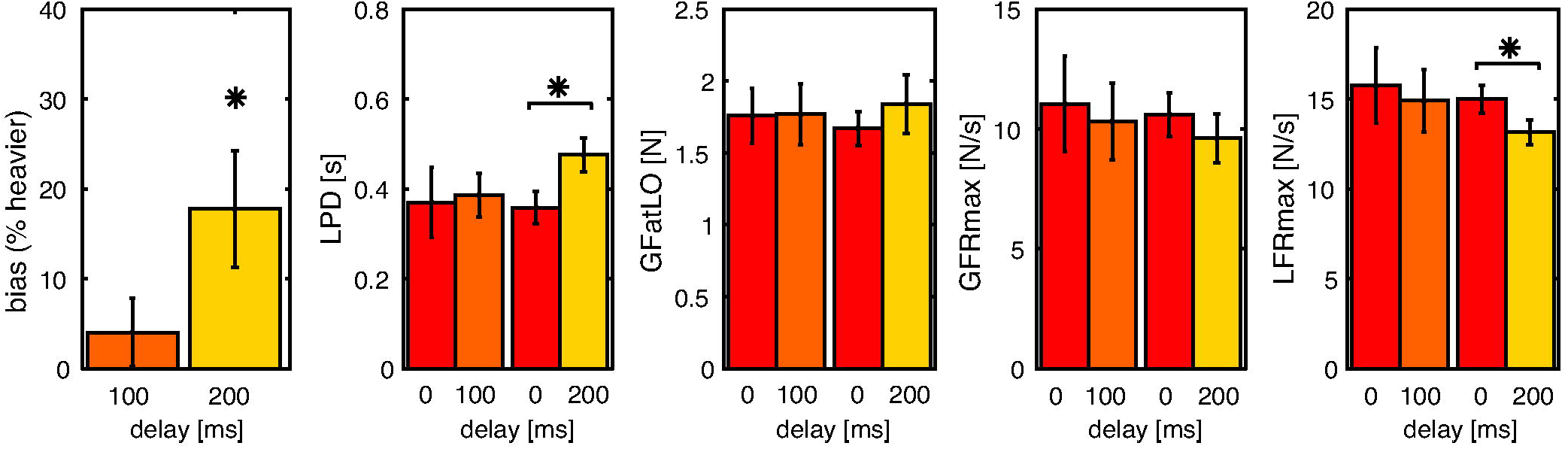
Results for Experiment 2. From left to right: perceptual biases, load phase duration (LPD), grip force at lift-off (GFatLO), maximum grip force (GFRmax) and load force rate (LFRmax). Lighter shades indicate larger delays. Error bars represent standard errors. Note that with 200 ms delay, there was a significant bias and significant difference in LPD and LFRmax (*p<0.05, paired samples t-test).

In addition, we examined force parameters in this experiment as well. As can be seen in Figure 5, differences between delay and no-delay lifts are more pronounced with a delay of 200 ms. Although GFatLO appears to increase with a delay, it was not significantly modified by the delay. No significant effect of delay was found for the GFRmax either. On the other hand, significant effects were found for LFRmax and LPD. LFRmax was lower with delay trials (*F*_(1,14)_=8.4, *p*=0.012, *η_p_*^2^=0.37). In accordance with the lower force rate, LPD was longer with a delay (*F*_(1,14)_=5.7, *p*=0.032, *η_p_*^2^=0.29). No effects of group or interactions of group × delay were found in any of the force parameters. Average force curves (not shown) were similarly changed as in Experiment 1. These effects were more pronounced for the 200 ms delay condition.

In sum, we found that a cube lifted with a visual delay could be perceived as 18% heavier compared to a cube lifted without such a delay. In addition, the slower force generation found in Experiment 1 was replicated in Experiment 2. To examine whether differences in force parameters were related to perceptual changes, the relative changes in delay trials vs. no delay trials were correlated with perceptual biases. For the 100 ms delay, a larger bias was found to correlate with an increase in LFRmax (*R*=0.85, *p*=0.007). In addition, a decrease in LPD was associated with a larger bias (*R*=−0.78 *p*=0.023). These results are unexpected, since on average smaller values of LFRmax and a larger LPD in combination with larger biases were found. No correlations of biases with GFRmax and GFatLO were found. None of the correlations between force parameters and perceptual biases were significant for the 200 ms conditions.

### Visual and haptic information both contribute independently to force scaling

Results from both experiments show that the force scaling is altered in response to a visual delay. This effect on force parameters in object lifting is seen very early in the force profile, suggesting that the object contact time point might be an important event. In general, force generation slows down, but it also appears that force rate profiles are shifted in time. To get a better insight into the mechanisms underlying these effects on force generation, we applied different models to the load force traces (Figure 2A), which tested how haptic and visual feedback was incorporated online into force generation, before object lift-off. We considered three models: a shift, stretch and sum model. In the shift model, only the onset of the force curve is shifted in time. In contrast, in the stretch model, the duration of force generation is increased. In the shift and stretch models, it is assumed visual and haptic feedback is combined before affecting the force curve. In the sum model, two force curves (one for each feedback modality) are formed and then combined. How much the non-delayed force curve is shifted, stretched or reshaped depends on the relative weight given to vision and haptics. In this way, we could estimate how visual and haptic information are integrated for force scaling during object lifting.

An example of three fitted curves to the no-delay and the 200 ms delay conditions of a representative participant lifting a 200 g cube is shown in Figure 2B. Average RMSE values are shown in Figure 6A. It can be seen that the sum model outperforms the other models in all conditions in both experiments. To test whether this was significant, a repeated measures ANOVA was performed with model as a factor. For Experiment 1, a 3 (model) × 2 (delay) × 4 (mass) ANOVA revealed effects of model (*F*_(2,26)_=3.9, *p*=0.034, *η_p_*^2^=0.23) and mass (*F*_(1.6,20.7)_=14.3, *p*<0.001, *η_p_*^2^=0.52). Post-hoc tests indicated that the sum model was significantly better than the stretch model. In addition, it was found that the models performed better for lower masses. An interaction between model × delay (*F*_(2,26)_=3.5, *p*=0.046, *η_p_*^2^=0.21) had no significant post-hoc tests. For Experiment 2, a 3 (model, within factor) × 2 (group, 100 ms and 200 ms, between factor) ANOVA showed no effect of model, group or an interaction effect.

**Figure 6.**
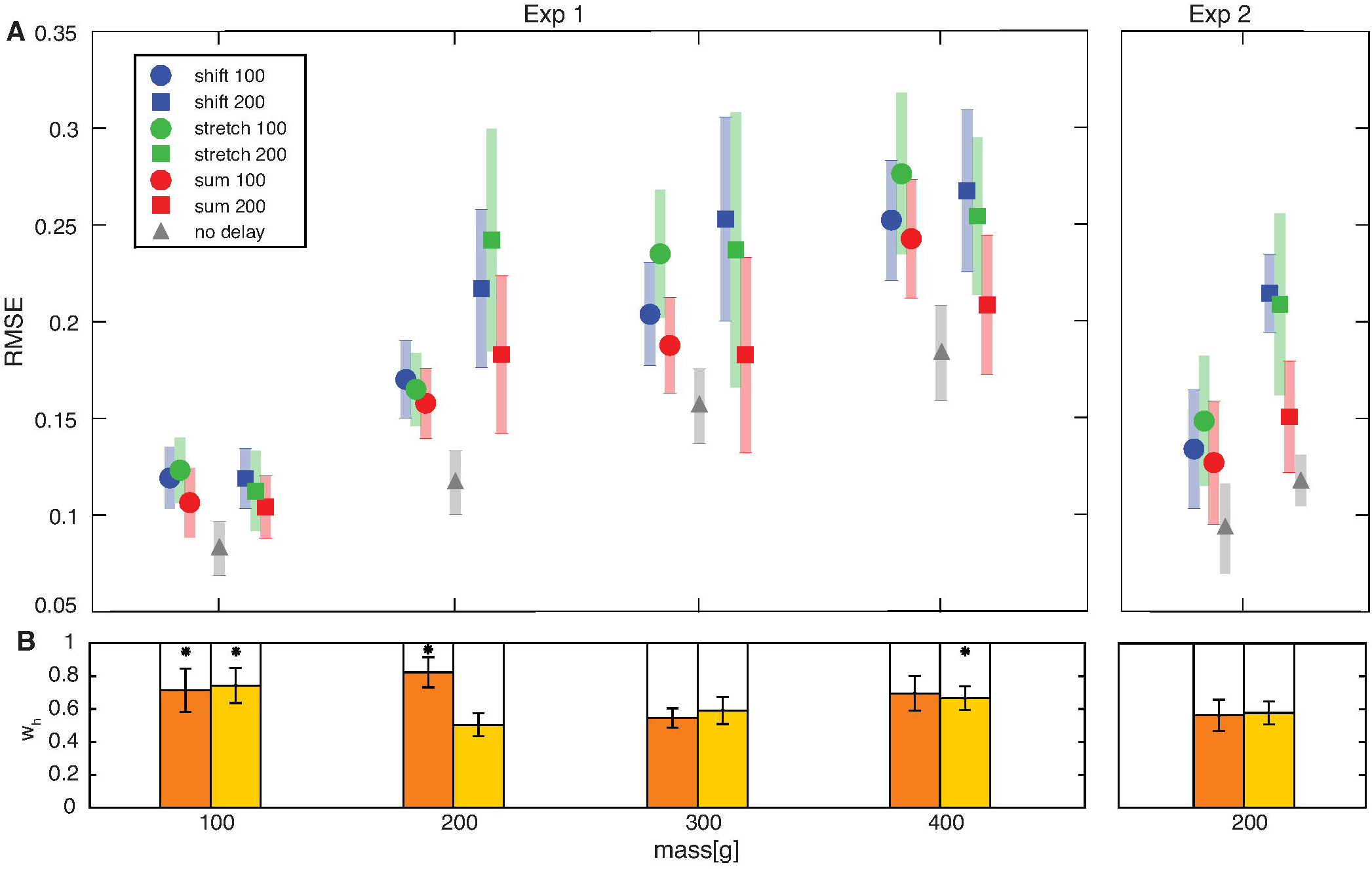
A. RSME for the shift (blue), stretch (green) and sum (red) model, for Experiment 1 (left panel) and Experiment 2 (right panel). Circles indicate 100 ms delay and squares 200 ms conditions. Grey triangles represent the no-delay (baseline) values. Two separate no-delay values are displayed in Experiment 2, one for each subject group. Shaded areas represent standard errors. Note that the sum model performs best in all conditions. B. Weights for the sum model for Experiment 1 (left panel) and Experiment 2 (right panel). Orange and yellow bars represent 100 ms and 200 ms delay conditions, respectively. Bars represent w_h_ (colored portion) and w_v_ (blank portion). Asterisks indicate significant difference from 0.5 (one sample t-test, p<0.05). Error bars represent standard errors.

The weights that were given to the haptic information (*w_h_*) and visual information (*w_v_*) for the sum model are shown in Figure 6B. It can be seen that most values are between 0 and 1, indicating that both visual and haptic information contribute to the modeled force curve. Indeed, all weights were significantly different from 0 and from 1 (one sample t-test). In some conditions, values were significantly higher than 0.5 (one sample t-test), indicating a larger weight for haptic information. The weights were not correlated with the magnitude estimations (z-scores, Experiment 1) or with the perceptual biases (Experiment 2). The average weight for vision over all conditions and experiments was 0.36.

## Discussion

The aim of this study was to investigate the relative weighting of visual and haptic information during object lifting. To be able to disentangle the contribution of visual and haptic signals, we used a virtual reality environment where vision could be desynchronized from haptics. We found that visual delay did not only alter the force scaling during lifting, but also increased the perceived heaviness of lifted objects. Our results indicate that, even though haptic information should be sufficient to lift an object skillfully after secure contact points have been made, visual inputs play a substantial role in controlling object lifting and heaviness perception. A comparison of different models explaining how multisensory information is integrated at discrete sensorimotor control points showed that visual and haptic feedback contribute rather independently to force scaling.

### Visual delay alters force scaling

We initially hypothesized that, with a visual delay, forces would continue to increase after haptic lift-off, because the object would not yet visibly move. However, differences between force profiles of delayed and non-delayed lifts were already seen soon after object contact. This suggests that object contact is an important event in object lifting. Although tactile mechanoreceptors in the fingertips sense contact with the object (Johansson and Flanagan 2009), the fingers do not yet touch the object visibly. This mismatch of expected contact might lead to uncertainty, slowing down the force generation profile. We found that the reaching phase was also affected by visual delay, indicating that a mismatch might already be perceived before contact. However, if participants were aware of this visuo-haptic asynchrony, they were not able to correct for it in their force scaling. Earlier studies also found effects of visual delay on reaching (Foulkes and Miall 2000; Shimada et al. 2004), suggesting that there is a general effect of visuo-haptic asynchronies in motor control. It is not obvious that reaching and lifting (or force control) is similarly affected by sensory information, as different responses have been shown as well (Brenner and Smeets 1996; Danion et al. 2013) and different parameters (e.g. finger positioning and fingertip forces) have to be controlled. Therefore, our findings extend the existing knowledge on motor control by showing how visuo-haptic integration is processed in a rapid succession of key sensorimotor control points, a feature of lifting movements. How similar this integration is for lifting kinetics vs reaching kinematics should be determined in future research.

At later stages, the force scaling was also affected, where a visual delay increased grip forces at lift-off. A second mismatch between haptic and visual lift-off might have caused the further increases in grip force after the object was lifted from the surface. The phantom devices do not provide similar tactile information about object friction and material as real objects would. This might explain why we have limited effects on grip force rates, which are naturally adjusted to both the weight and friction of objects (Johansson and Westling 1984).

### Visual delay makes lifted objects feel heavier

A second main finding of our study was that cubes lifted with a visual delay were perceived as being heavier than cubes lifted without a delay. When cubes were compared directly, the cube lifted with delayed vision was perceived as 18% heavier (= a perceptual effect of 36 g for a 200 g object lifted with a 200 ms visual delay, Experiment 2). These results extend similar findings on compliance perception (Di Luca et al. 2011) and heaviness perception in object transport (Honda et al. 2013) to the complete action of lifting and holding objects.

The altered force profiles following visual delay show similarities to force scaling of lifted objects that are heavier than expected (Johansson and Westling 1988), suggesting an increased sense of effort (van Polanen and Davare 2015). On the other hand, forces and sense of effort are not always related to heaviness perception, as seen in the size-weight illusion that occurs independent of force alterations (Flanagan and Beltzner 2000; Grandy and Westwood 2006). Although we found both effects on heaviness perception and force scaling, these effects were not strongly correlated. Therefore, it seems likely that the increased heaviness perception is not entirely caused by the altered lifting kinematics, and might be an independent process. In a previous experiment where objects needed to be lifted and rated for their heaviness, we found a relation between grip force rate and weight perception (van Polanen and Davare 2015). In the present experiment, there were no effects found on grip force rate, which could explain the different results in both studies. It is possible that an increased sense of effort is more apparent when changes in grip forces occur, as also seen in other studies (Flanagan and Bandomir 2000; Flanagan et al. 1995). In the virtual reality, grip forces might be less accurately scaled, since feedback about friction cues is absent. Moreover, the current study design essentially generates a mismatch in sensory feedback loops, whereas in the former study (van Polanen and Davare 2015) effects were driven by manipulations of predictive mechanisms (i.e. expected force scaling based on predicted object weight). Furthermore, although the loading phase seems important for weight perception, visuo-haptic asynchronies in the phases after lift-off could affect the weight perception as well, such as the transitional phase or the replacement of the object on the table. Kinematic visual information has been shown to provide information about object weight (Hamilton et al. 2007; Runeson and Frykholm 1981; Streit et al. 2007). As we stated in the Introduction, multisensory integration for perception might be different than for action control, because for perception, sensory inputs can be acquired over longer time periods whereas brief key events are important for action control. Therefore, the brain processing or weighting of the visual and haptic modalities might have differed between heaviness perception and force scaling, resulting in the absence of strong correlations. To summarize, although perceptual and force scaling effects were observed in similar directions, they do not seem to be caused by the same underlying mechanisms.

### Visual and haptic information are independently weighted to control force scaling

To further investigate the integration of visual and haptic feedback for force scaling, we compared different models in which we examined how these sensory inputs are weighted. Our comparisons rule out two models of force control during object lifting. First, the onset of force generation does not seem to be delayed until an integrated feedback of visual and haptic contact can be formed, which rules out the shift model. Secondly, visual delay does not simply slow down force generation until an integrated percept of lift-off is determined, which rules out the stretch model. In contrast, it appears that both feedback of haptic and visual object contacts trigger changes in force generation and both modalities contribute to the final force output. Interestingly, this suggests that haptic and visual information is not ‘merged’ to control force scaling (e.g. to initiate force build-up or determine its duration in the shift and stretch model, respectively) but both influence force control individually. The notion that multimodal information is not merged agrees with earlier research in size perception that showed that information from individual senses is not lost in the perception of a single object (Hillis et al. 2002) and not completely fused in sequential event detection (Bresciani et al. 2006). The largest effects of visual delay on the force generation profiles are seen if equal weighting is assigned to visual and haptic feedback, leading to lower load force rates and longer loading phase durations, exactly as seen in our experimental data. The average weights given to haptic and visual information from our model fits suggest that while haptic feedback has a greater influence on the control of object lifting, visual signals play a substantial role in shaping up the motor commands.

A limitation of the present model is that it only considers a preplanned bell-shaped force output, starting from visual and/or haptic object contact. That is, corrective feedback processes that could be involved after lift-off are not included in the model. Another issue is that although the sum model outperformed the shift and stretch models, this does not necessarily indicate this model fits the data optimally. We used root mean square errors to provide an estimation of goodness of fit and discrepancies between the data and the model remain, even in the sum model. Since force scaling in object lifting is a complex process, which requires tight coordination between two fingers and multiple muscles, it seems obvious that more complicated models might capture force data more optimally. However, it was beyond the scope of this paper to model these processes. Our models provide new insight into how visual and haptic feedback about object contact is integrated and used to update a motor plan online.

Although in general we found that force scaling and weight perception is influenced by visual and haptic information, the actual values (e.g. model weights and biases) differed among participants. These results suggest that the relative reliance on vision differs among individuals. Possibly, participants who are more skilled in fine motor control or are better at haptic sensing might give a bigger weight to haptic information. Furthermore, participants might shift their weighting if they are more familiar with virtual environments. It needs to be determined whether these weightings are constant for a specific individual over time.

Another relevant question is why we would rely on visual information at all? We are perfectly able to lift objects with our eyes closed, without any visual information. Studies on the integration of haptic and visual information in perception suggest that this integration is an automatic process (Helbig and Ernst 2008). Furthermore, the visual contribution can even be higher than optimal predictions, e.g. in compliance perception (Cellini et al. 2013) and torque perception (Xu et al. 2012). Although there is evidence that the relative weighting of haptic and visual information can be adapted to the task (Takahashi and Watt 2014), the shift to a particular modality might not always be complete (Drewing and Kruse 2014). Overall, this literature suggests that visual information will be used when present, even if not necessarily needed. However, this research about haptic and visual integration concerns perceptual measures, so this might not hold true for sensorimotor tasks.

It is possible that our delay manipulation in the virtual reality slightly altered the natural weighting of haptic and visual information occurring when lifting real objects. However, it is unlikely that our findings in virtual reality are completely different from visuo-haptic integration in real life conditions. First, given that delay and no-delay trials were presented randomly from trial to trial, participants could not adapt to visual delay and thus change their motor control strategies. This is also in line with the fact that the weighting of visual and haptic information cannot be changed flexibly (Cellini et al. 2013). Secondly, since we performed our control condition (no-delay) and our manipulation (delay) both in virtual reality, the comparison between them still informs about how multisensory information is processed by the brain. Any changes in signal accuracy due to the virtual environment that could affect the weighting between vision and haptics (Ernst and Banks 2002) would be present in both conditions. Thirdly, possible general changes in lifting behavior, e.g. less skilled lifting due to the absence of frictional cues, were present in lifts with and without delay and would not explain our effects of delay.

### Concluding comments

In sum, our results provide the first account for the organization of multisensory integration when action control is performed in a rapid series of sensorimotor control points, such as in lifting movements. This experiment was performed in virtual reality and might contribute to developing virtual environments where both visual and haptic feedback is vividly rendered. For instance, in depth measurements could improve research into the influence of a visual delay in robotic surgery (Kim et al. 2005; Onda et al. 2010). On the other hand, our findings also open up the possibility to create an illusionary heavy object simply by increasing the visual delay.

## Supporting information

Supplemental model code

## Acknowledgements

VVP was funded by a Fonds Wetenschappelijk Onderzoek grant (Post-doctoral fellowship, FWO, Belgium, 12X7118N). MD was funded by a Fonds Wetenschappelijk Onderzoek grant (FWO Odysseus, Belgium FWO: G/0C51/13N). RT and MD were funded by a BBSRC David Phillips fellowship (UK) to MD (BBSRC: BB/J014184/1). AN was funded by a JSPS KAKENHI Grant (JP23700608). The authors would like to thank Dr. Atsuo Nuruki and Dr. Raz Leib for their help in designing the virtual environment.

## References

Brenner E, and Smeets JBJ. Size illusion influences how we lift but not how we grasp an object. Experimental brain research 111: 473–476, 1996.

Bresciani J-P, Dammeier F, and Ernst MO. Vision and touch are automatically integrated for the perception of sequences of events. Journal of vision 6: 2–2, 2006.

Buckingham G, and Goodale MA. Lifting without seeing: the role of vision in perceiving and acting upon the size weight illusion. PloS one 5: e9709, 2010.

Buckingham G, Wong JD, Tang M, Gribble PL, and Goodale MA. Observing object lifting errors modulates cortico-spinal excitability and improves object lifting performance. Cortex; a journal devoted to the study of the nervous system and behavior 50: 115–124, 2014.

Cellini C, Kaim L, and Drewing K. Visual and haptic integration in the estimation of softness of deformable objects. i-Perception 4: 516–531, 2013.

Danion F, Diamond JS, and Flanagan JR. Separate Contributions of Kinematic and Kinetic Errors to Trajectory and Grip Force Adaptation When Transporting Novel Hand-Held Loads. Journal of Neuroscience 33: 2229–2236, 2013.

Di Luca M, Knörlein B, Ernst MO, and Harders M. Effects of visual-haptic asynchronies and loading-unloading movements on compliance perception. Brain Research Bulletin 85: 245–259, 2011.

Drewing K, and Kruse O. Weights in Visuo-Haptic Softness Perception are not Sticky. In: EuroHaptics 2014, Part I, LNCS 8618, edited by Auvray M, and Duriez C. Berlin Heidelberg: Springer-Verlag, 2014, p. 68–76.

Ernst MO, and Banks MS. Humans integrate visual and haptic information in a statistically optimal fashion. Nature 415: 429–433, 2002.

Flanagan JR, and Bandomir CA. Coming to grips with weight perception: effects of grasp configuration on perceived heaviness. Percept Psychophys 62: 1204–1219, 2000.

Flanagan JR, and Beltzner MA. Independence of perceptual and sensorimotor predictions in the size-weight illusion. Nature neuroscience 3: 737–741, 2000.

Flanagan JR, Wing AM, Allison S, and Spenceley A. Effects of Surface Texture on Weight Perception When Lifting Objects with a Precision Grip. Percept Psychophys 57: 282–290, 1995.

Foulkes AJM, and Miall RC. Adaptation to visual feedback delays in a human manual tracking task. Experimental brain research 131: 101–110, 2000.

Gibo TL, Bastian AJ, and Okamura AM. Grip force control during virtual object interaction: effect of force feedback,accuracy demands, and training. IEEE transactions on haptics 7: 37–47, 2014.

Grandy MS, and Westwood DA. Opposite perceptual and sensorimotor responses to a size-weight illusion. Journal of neurophysiology 95: 3887–3892, 2006.

Hamilton AF, Joyce DW, Flanagan JR, Frith CD, and Wolpert DM. Kinematic cues in perceptual weight judgement and their origins in box lifting. Psychol Res 71: 13–21, 2007.

Helbig HB, and Ernst MO. Optimal integration of shape information from vision and touch. Experimental brain research 179: 595–606, 2007.

Helbig HB, and Ernst MO. Visual-haptic cue weighting is independent of modality-specific attention. Journal of vision 8: 1–16, 2008.

Hillis JM, Ernst MO, Banks MS, and Landy MS. Combining Sensory Information: Mandatory Fusion Within, but Not Between, Senses. Science 298: 1627–1630, 2002.

Honda T, Hagura N, Yoshioka T, and Imamizu H. Imposed visual feedback delay of an action changes mass perception based on the sensory prediction error. Frontiers in psychology 4: 760, 2013.

Johansson RS, and Flanagan JR. Coding and use of tactile signals from the fingertips in object manipulation tasks. Nature reviews Neuroscience 10: 345–359, 2009.

Johansson RS, and Westling G. Coordinated isometric muscle commands adequately and erroneously programmed for the weight during lifting task with precision grip. Experimental brain research 71: 59–71, 1988.

Johansson RS, and Westling G. Roles of Glabrous Skin Receptors and Sensorimotor Memory in Automatic-Control of Precision Grip When Lifting Rougher or More Slippery Objects. Experimental brain research 56: 550–564, 1984.

Kambara H, Shin D, Kawase T, Yoshimura N, Akahane K, Sato M, and Koike Y. The effect of temporal perception on weight perception. Frontiers in psychology 4: 40, 2013.

Kim T, Zimmerman PM, Wade MJ, and Weiss CA. The effect of delayed visual feedback on telerobotic surgery. Surgical Endoscopy and Other Interventional Techniques 19: 683–686, 2005.

Knill DC, and Saunders JA. Do humans optimally integrate stereo and texture information for judgments of surface slant? Vision research 43: 2539–2558, 2003.

Monzée J, Lamarre Y, and Smith AM. The effects of digital anesthesia on force control using a precision grip. Journal of neurophysiology 89: 672–683, 2003.

Onda K, Osa T, Sugita N, Hashizume M, and Mitsuishi M. Asynchronous force and visual feedback in teleoperative laparoscopic surgical system. IEEE/RSJ 2010 International Conference on Intelligent Robots and Systems, IROS 2010 - Conference Proceedings 844–849, 2010.

Runeson S, and Frykholm G. Visual perception of lifted weight. Journal of experimental psychology Human perception and performance 7: 733–740, 1981.

Sarlegna FR, Baud-Bovy G, and Danion F. Delayed visual feedback affects both manual tracking and grip force control when transporting a handheld object. Journal of neurophysiology 104: 641–653, 2010.

Shimada S, Hiraki K, Matsuda G, and Oda I. Decrease in prefrontal hemoglobin oxygenation during reaching tasks with delayed visual feedback: a near-infrared spectroscopy study. Cognitive Brain Research 20: 480–490, 2004.

Streit M, Shockley K, Riley MA, and Morris AW. Rotational kinematics influence multimodal perception of heaviness. Psychonomic Bulletin & Review 14: 363–367, 2007.

Takahashi C, and Watt SJ. Visual-haptic integration with pliers and tongs: Signal “weights” take account of changes in haptic sensitivity caused by different tools. Frontiers in psychology 5: 1–14, 2014.

Uçar E, and Wenderoth N. Movement observation affects sensorimotor memory when lifting a familiar object. Cortex; a journal devoted to the study of the nervous system and behavior 48: 638–640, 2012.

van Polanen V, and Davare M. Sensorimotor Memory Biases Weight Perception During Object Lifting. Frontiers in human neuroscience 9: 700, 2015.

Xu Y, O’Keefe S, Suzuki S, and Franconeri SL. Visual influence on haptic torque perception. Perception 41: 862–870, 2012.

Zwislocki JJ, and Goodman DA. Absolute scaling of sensory magnitudes: a validation. Percept Psychophys 28: 28–38, 1980.

