## Supplemental model code for "Visual delay affects force scaling and weight perception when lifting objects in virtual reality"

### Main function

```
function [rmse,w_h]=model_main3(force_nd,force_d,obj_weight,delay)

% input:
% force_nd      data of no-delay condition (1 subject)
% force_d       data of delay condition (1 subject)
% obj_weight    object mass (N)
% delay         visual delay (s)
%
% output:
% rmse          root mean square error between model and data
% w_h           weight of haptic feedback

% initialize
w_h=ones(1,3);
rmse=zeros(1,3);

t0=0; % time of object contact
sfs=500; % sample frequency data

% Fit duration d on no-delay data
% duration boundaries are set between 0 and 2s.
d=fminbnd(@(d) modelsinefun(d,...
        obj_weight,delay,t0,force_nd,sfs,1,1),0,2);


% Fit w_h for each model on delay data by minimizing rmse
% w_h boundaries are set between 0 and 1
for mtype=1:3
    w_h(mtype)=fminbnd(@(w) modelsinefun(d,...
        obj_weight,delay,t0,force_d,sfs,w,mtype),0,1);

    rmse(mtype)=...
        modelsinefun(d,obj_weight,delay,t0,...
        force_d,sfs,w_h(mtype),mtype);
end
```

### Model and fit function

```
function
 rmse=modelsinefun(d,obj_weight,delay,t0,force_data,sfs,w_h,mtype)

% input:
% d             planned duration (in s)
% obj_weight    weight object (in N)
% delay         visual delay (in s)
% t0            time of object contact - GF onset (in s)
% force_data    measured force data (in N)
% sfs           sample frequency of measured force
% w_h           weight of haptic feedback
% mtype         model type: 1) summation 2) move 3) stretch
```

```
%
% output:
% rmse          root mean square error between model and data
%
% 1 - sine summation
%   two acceleration sines that are summed
%   Visual sine is displayed with respect to haptic sine with delay
%
% 2 - sine shift
%   one acceleration sine that is shifted by a weighted delay period
%
% 3 - sine stretch
%   one acceleration sine that is stretched by a weighted duration
%   time


fs=10000;
maxtijd=2.5;

liftoff=find(force_data>obj_weight,1);

% comparision ranges
samplerangedata=fix(1:liftoff);
samplerangemodel=fix(1:fs/sfs:liftoff*fs/sfs);

% weight of vision
w_v=1-w_h;

% new duration definition
if mtype==3 % model 3: stretch
    dnew=d+w_v*delay;
else % model 1/2: sum/shift
    dnew=d;
end

% time steps
t=[t0;t0+dnew/2;t0+dnew]*fs;
if mtype==2 % model 2: shift
    t(:,2)=t(:,1)+w_v*delay*fs;
else % model 1/3: sum/stretch
    t(:,2)=t(:,1)+delay*fs;
end
t=fix(t);

acc_i=zeros(fs*maxtijd,2);

FR=obj_weight/dnew;
G=2*pi*FR/dnew;

% calculate sine
if mtype==1 % model 1: sum
    acc_i(t(1,1):t(3,1),1)=...
        G*w_h*sin((0:t(3,1)-t(1,1))*2*pi/(dnew*fs));
    acc_i(t(1,2):t(3,2),2)=...
```

```
        G*w_v*sin((0:t(3,2)-t(1,2))*2*pi/(dnew*fs));
elseif mtype==2 % model 2: shift
    acc_i(t(1,2):t(3,2),2)=...
        G*sin((0:t(3,2)-t(1,2))*2*pi/(dnew*fs));
else % model 3: stretch
    acc_i(t(1,1):t(3,1),1)=...
        G*sin((0:t(3,1)-t(1,1))*2*pi/(dnew*fs));
end

% integrate
acc=sum(acc_i,2);
vel=cumsum(acc)/fs;
forcesim=cumsum(vel)/fs;

% rmse
rmse=sqrt(mean((forcesim(samplerangemodel)-...
    force_data(samplerangedata)).^2));

end

end
```

*Published with MATLAB® R2017b*
